# Microtubules under mechanical pressure can breach dense actin networks

**DOI:** 10.1101/2023.05.15.540482

**Authors:** Matthieu Gélin, Alexandre Schaeffer, Jérémie Gaillard, Christophe Guérin, Benoit Vianay, Magali Orhant-Prioux, Marcus Braun, Christophe Leterrier, Laurent Blanchoin, Manuel Théry

## Abstract

The crosstalk between actin network and microtubules is key to the establishment of cell polarity. It ensures that the asymmetry of actin architec ture along cell periphery directs the organization of microtubules in cell interior. In particular, the way the two networks are physically intertwined regulates the spatial organization and the distribution of forces in the microtubule network. While their biochemical crosstalk is getting uncovered, their mechanical crosstalk is still poorly understood. Here we designed an in vitro reconstitution assay to study the physical interaction between dynamic microtubules with various structures made of actin fil aments. We found that microtubules can align and move by their polymerization force along linear bundles of actin filaments. But they cannot enter dense and branched actin meshworks, such as those found in the lamellipodium along cell periphery. However, when microtubules are immobilized, by their crosslinking with actin structures or others means, the force of polymerization builds up pressure in the microtubules that is sufficient to allow them to breach and penetrate these dense actin meshworks. This mechanism may explain the final progression of microtubules up to cell periphery through the denser parts of the actin network.

## Introduction

Cell polarity and internal compartmentalization depend on the interplay between actin filaments and microtubules which are both responsible for the positioning of organelles and the organization of endomembrane networks (Bornens, 2008; Akhmanova and Kapitein, 2022). Defective coupling between actin filaments and microtubules perturb cell ability to sense and respond to extracellular signals due to misoriented polarity, growth and migration (Ballestrem et al., 2000; Wu et al., 2008; Zhou et al., 2002). It is thus key to understand the rules regulating the mechanical and biochemical interaction of actin filaments and microtubules. While some shared signalling pathways have been shown to be key for their crosstalk (Krendel et al., 2002; Rooney et al., 2010), the process regulating their mechanical interaction is still unclear (Dogterom and Koenderink, 2019).

The actin network lies along cell periphery. In response to adhesive cues, actin filaments can self-organize into various structures (Blanchoin et al., 2014). Cell adhesion promotes the polymerization of actin filaments and their assembly into dense and branched meshworks, which can push and deform the plasma membrane (DeMali et al., 2002; Swaminathan et al., 2016). Above non-adhesive regions, the network contraction by myosins led to the coalescence of filaments into tight bundles connecting cell anchorage sites (Vignaud et al., 2020). Thus, this self-organization process somehow translates adhesive cues into intracellular actin organization and map in the cell the topology of the extracellular environment. This internal pattern of actin structures further serves as grounding for the self-organization of other cell cytoskeleton networks, which adapt their architecture to these guiding cues and thereby ensure the consistency of the entire cellular organization with the configuration of the extra-cellular environment (Bornens, 2008).

Microtubules respond specifically to different types of actin network architectures. They were described to grow along linear bundles of actin filaments (Burnette et al., 2007; Wu et al., 2008) but to be blocked by dense and branched actin meshwork in the lamellipodium (Ballestrem et al., 2000; Waterman-Storer and Salmon, 1997; Dema et al., 2023). Actin inward flow was shown to push, reorient and even break microtubules (Gupton et al., 2002; Ning et al., 2016; Salmon et al., 2002; Schaefer et al., 2002). However, microtubules penetration through actin meshworks has been observed in neuronal growth cone (Burnette et al., 2007) and at the front of migrating epithelial cells (Wittmann et al., 2003). On one hand the frequency of these “pioneer” or “exploratory” microtubules was increased by actin disassembly (Burnette et al., 2007) but on the other it was reduced by Rac inactivation (Wittmann et al., 2003). So the rules regulating MT stumbling against or breaching the dense actin meshwork in the lamellipodium along cell periphery are unclear. Nonetheless, it is key to understand this intertwining of the two networks since it regulates the balance of forces in the microtubule network and the orientation of cell polarity (Burute et al., 2017; Jimenez et al., 2021; Pitaval et al., 2017; Yamamoto et al., 2022) as well as the induction of membrane protrusion and directionality of cell migration (Schaefer et al., 2008; Dema et al., 2023; Bouchet et al., 2016; Bouchet and Akhmanova, 2017).

## Results

### Microtubule penetration in the lamellipodium of cultured cells

By looking at microtubule organizations in mouse embryonic fibroblasts, we could clearly see microtubules alignment along stress fibers (Figure 1A). Interestingly, these microtubules could reach the cell periphery in regions devoid of lamellipodium-like meshwork (Figure 1A’) but were stopped by such structure in others (Figure 1A”). Similarly, in fibroblast-like cells COS-7, MTs could barely progress up to cell periphery because of the dense actin meshwork in the lamellipodium (Figure 1B). However, few microtubules managed to pass through (Figure 1B’) and this was systematic at tip of membrane protrusions (Figure 1B’). This clear correlation suggested that either the rare and random penetration of microtubules promote actin reorganisation in the lamellipodium and formation of protrusions, or that the local organisation of actin filaments that can promote the formation of protrusion is permissive to microtubule penetration. We could document these rare events of microtubule piercing the lamellipodium of live epithelial-like cells, PtK_2_ (Figure 1C, top. Supplementary Figure S1) expressing tubulin-GFP and labelled with SiR-actin. However, we could not identify any characteristic local feature of the actin network where microtubule breached through (Figure 1C, bottom. Supplementary Figure S1). In some instances, actin network was quite homogeneous (Figure 1C Left) whereas is others microtubules seemed to grow along preexisting bundles of actin filaments (right). Since our ability to finely modulate the density and inner architecture of the lamellipodium in living cells is limited, we designed a dedicated reconstitution assay in vitro in order to test the role of actin network architecture to microtubule penetration.

**Figure 1.**
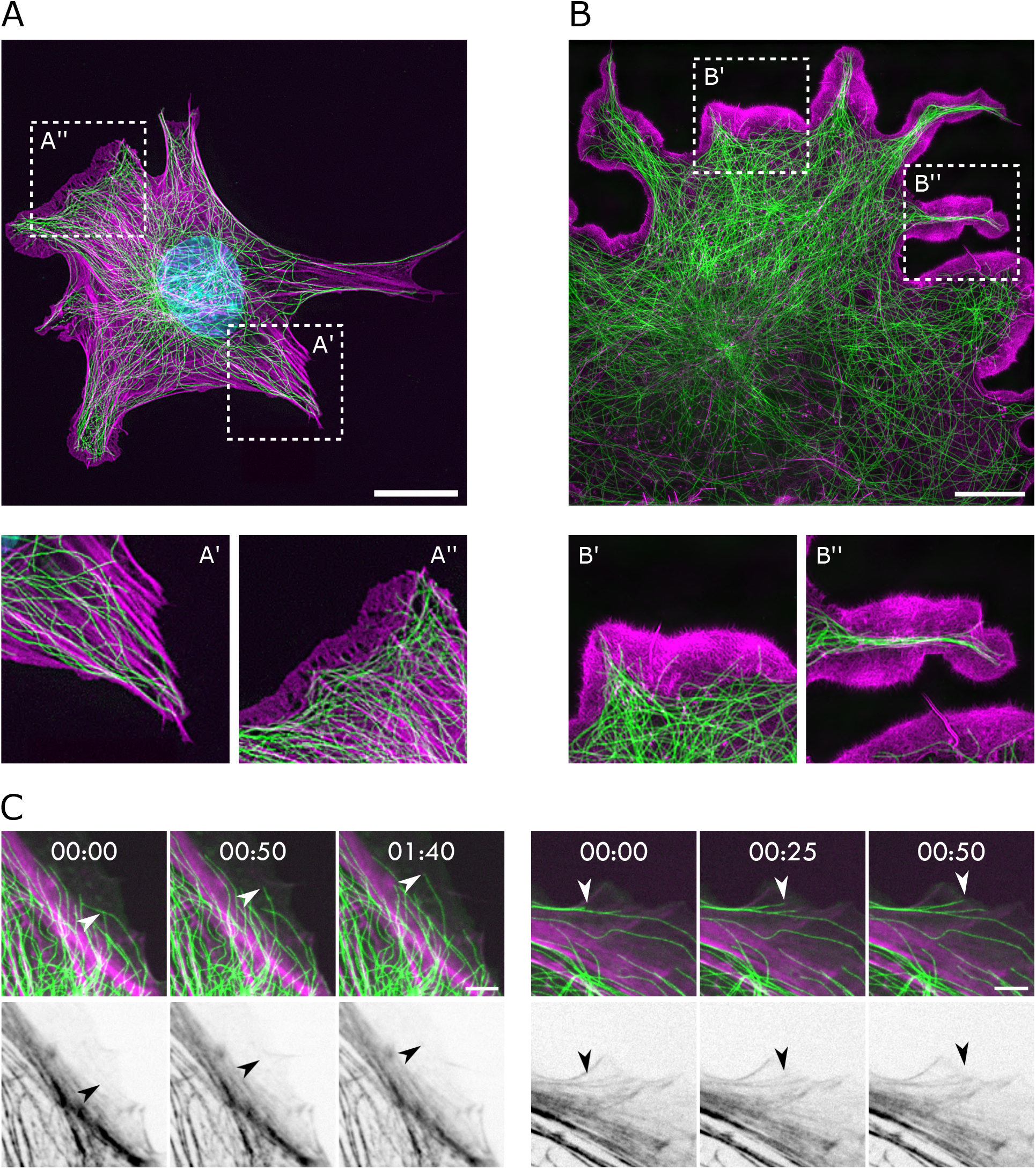
Microtubule actin interaction is structure dependent. (**A**) Immunofluorescence images of the actin (magenta) and microtubule (green) network found in a MEF cell, nucleus is in blue. Maximum projection, processing: gamma and unsharp mask. Scale bar, 20 µm. (**B**) Immunofluorescence images of the actin (magenta) and microtubule (green) network found in a COS-7 cell obtained by Structured Illumination Microscopy (SIM). Scale bar, 20 µm. (**C**) Time-lapses of PtK_2_ cells tubulin-GFP (green) incubated with 200nM of SiR-actin (magenta). The white arrows are showing microtubules entering the lamellipodial region of the cell (Top) and the corresponding black arrows are showing the actin structure encountered along their path (bottom). Scale bar, 5 µm.

Indeed, reconstitution assays offer the possibility to work with controlled and reproducible cytoskeleton networks. So far, they revealed that linear actin filaments can align with and orient microtubules (Farhadi et al., 2018; Lopez and Valentine, 2016; López et al., 2014; Kučera et al., 2022), and that branched actin filaments can limit microtubule mobility (Farhadi et al., 2020; Kučera et al., 2022) and block their elongation (Colin et al., 2018; Inoue et al., 2019), but these studies could not specifically address the mechanism of the piercing of dynamic microtubules into dense actin networks, which can occur in cells. This requires that microtubules and actin filaments are not randomly mixed in order for microtubules to first grow freely and then encounter dense actin meshwork. To that end, the two networks need to be spatially separated. In addition, microtubules should not be attached to the substrate so that they could move and deform freely as they encounter the actin network.

### Using lipid patterning to control actin architecture

To study the interaction between dynamic microtubules and a branched actin network, we first needed a way to control the architecture of this network in a delimited space. We first used the classical approach of grafting Nucleation Promoting Factor (NPF) on micropatterned regions on glass, as was done previously (Reymann et al., 2010). Coupling that with a control of actin elongation with Capping Protein (CP) (Boujemaa-Paterski et al., 2017), we obtained two types of actin architecture. One where the branched actin meshwork polymerized without CP and bundles could grow outside the pattern region (Figure 2B left). Another case where the branched actin network polymerizes in the presence of CP, which limits the growth of actin bundles (Figure 2B right). However, the use of CP led to an irreproducibility in the growth of the meshwork. In the examples shown in Figure 2C, the four images were taken near to each other on the same slide, yet networks displayed quite distinct densities and various internal level of heterogeneity. In an attempt to solve this reproducibility issue, we used micropatterned lipid bilayer, on which a Nucleation Promoting Factor (Snap-Streptavidin-WA-His) was attached via a biotin-streptavidin link. Indeed, using NPF grafted onto a lipid bilayer could increase by a factor of 10 the efficiency to generate actin assembly compared with NPF grafted onto a glass micropattern (Colin et al., 2022). Unexpectedly, this method unlocked a new level of control over the branched meshwork architecture. Indeed, we found that we could alter the architecture of the meshwork simply by changing the concentration of the Arp2/3 complex and without using CP (Figures 2E and F). At a low concentration of Arp2/3 complex (10nM), actin filaments could grow out of the branched meshwork and form bundles. In contrast, at a higher concentration of the Arp2/3 complex (100nM), branches were much more frequent and filaments were so numerous that they used up all the monomers, so that no filaments could grow out of the network and no outer bundles could form (Figure 2E). In addition, the use of a lipid bilayer also solved the reproducibility issue: as the actin network was nucleated much more rapidly all over the micropatterned area, the subsequent elongation of filaments led to quite homogeneous network over the stretch of several hundreds of microns (Figure 2F). These two conditions allowed us to test two characteristic architectures present at the cell periphery, namely dense networks of branched filaments and less dense networks with long bundles of linear filaments, and to study their impact on the behavior of dynamic microtubules.

**Figure 2.**
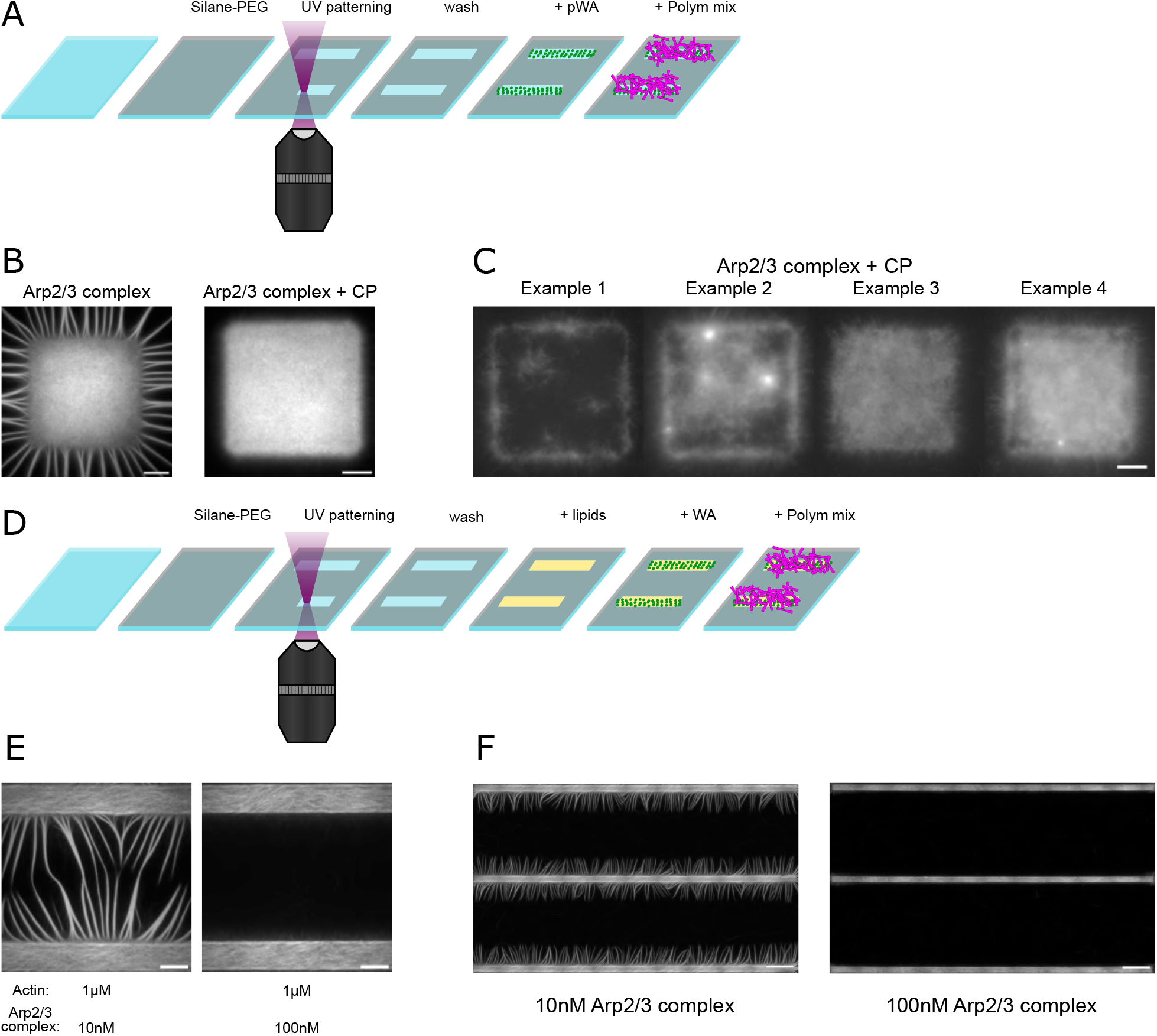
Using lipid patterning to control actin architecture. (**A**) Scheme of the method used to pattern NPFs on glass with the PRIMO micropatterning technique. (**B**) Images of branched actin network polymerized from a solid patterns in presence of 100nM of the Arp2/3 complex (left) or 100nM of the Arp2/3 complex and 3nM of Capping Protein (right). Scale bar, 10 µm. (**C**) Images of branched actin network polymerized from a solid patterns in presence of 100nM of the Arp2/3 complex and 3nM of Capping Protein. Scale bar, 10 µm. (**D**) Scheme of the method used to pattern lipids on which to graft NPFs with the PRIMO micropatterning technique. (**E**) Images of branched actin network polymerized from bar lipid patterns in presence of 10nM (left) or 100nM of the Arp2/3 complex (right). Scale bar, 10 µm. (**F**) Images of branched actin network polymerized from bar lipid patterns in presence of 10nM (left) or 100nM of the Arp2/3 complex (right). Scale bar, 50 µm.

### Alignment of dynamic microtubule along actin bundles

We then introduced free tubulin and microtubule seeds in the polymerization mix to study the interaction of dynamic microtubules with actin networks of different architectures. Microtubules exhibited normal behavior, growing and shrinking with their classical dynamic instability (Figure 3A). Surprisingly, when actin bundles were present (10nM of Arp2/3 complex), we observed an alignment between the microtubules and the bundles (Figure 3B). This observation was unexpected, as no actin-microtubule crosslinkers was present in the mix, contrary to previous studies (Lopez and Valentine, 2016; Willige et al., 2019). The effect was quite strong and concerned the majority of the microtubules (Figure 3C). However, since no crosslinker was present, the microtubules were free to diffuse along the length of the actin bundles, as is seen in figure 3B. Importantly, the presence or absence of actin bundles had no impact on the microtubule dynamics and thus on microtubule length distribution (Figures 3D, E).

**Figure 3.**
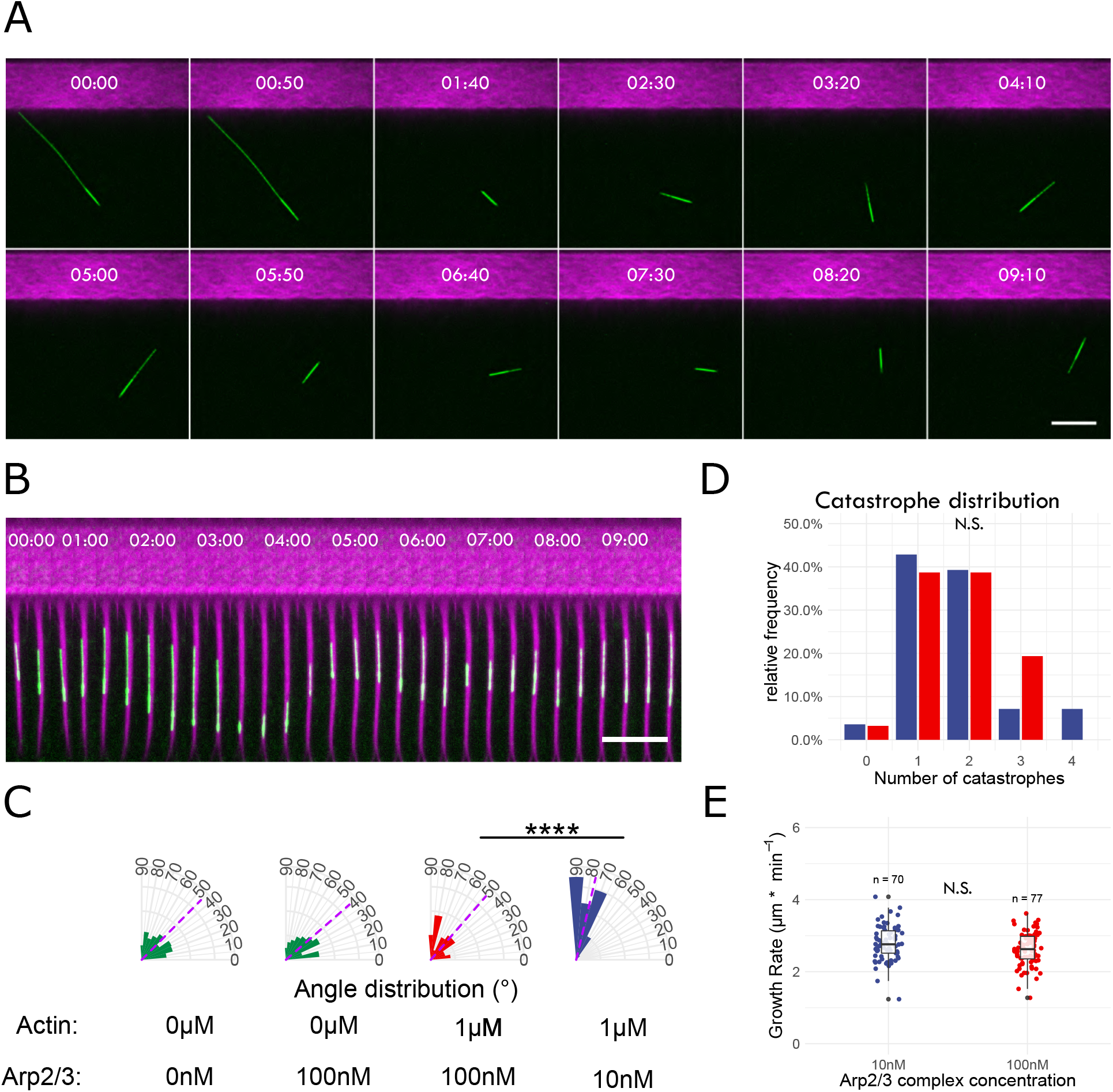
Microtubule alignment along actin bundles. (**A-B**) Time-lapse imaging of dynamic microtubules (green) polymerizing next to branched actin network (magenta) in absence (100nM of Arp2/3 complex, A) or presence (10nM of Arp2/3 complex, B) of actin bundles. (**C**) Distribution of microtubule orientation relative to the branched actin pattern. The magenta line correspond to the mean of the distribution. N.S. p>0.05, **** p<0.0001 (Student T-test). (**D**) Catastrophe distribution of microtubule interacting with the actin network in the presence (10nM of Arp2/3 complex, blue) or absence (100nM of Arp2/3 complex, red) of actin bundles. NS p>0.05 (Fisher test). (**E**) Growth rate distribution of microtubule interacting with the actin network in the presence (10nM of Arp2/3 complex, blue) or absence (100nM of Arp2/3 complex, red) of actin bundles. N.S. p>0.05 (Student T-test). Scale bar, 10 µm.

### Microtubule can breach the branched actin network

Interestingly, as microtubules growing tip encountered the branched actin network, the force produced by their polymerization, likely by a Brownian ratchet-like mechanism (Mogilner and Oster, 2003), was strong enough to propel them backward relative to the actin network (Figure 4A-B). Such a movement could be observed in the presence or absence of bundles, suggesting that the friction along the bundle was not sufficient to counteract the polymerization force (Figure 4A-B). Noteworthy, in our conditions the contact of the microtubule growing end with the actin network did not induce microtubule catastrophe (Janson et al., 2003) because the microtubule was free to move backward so the elongation of protofilament was not impaired.

**Figure 4.**
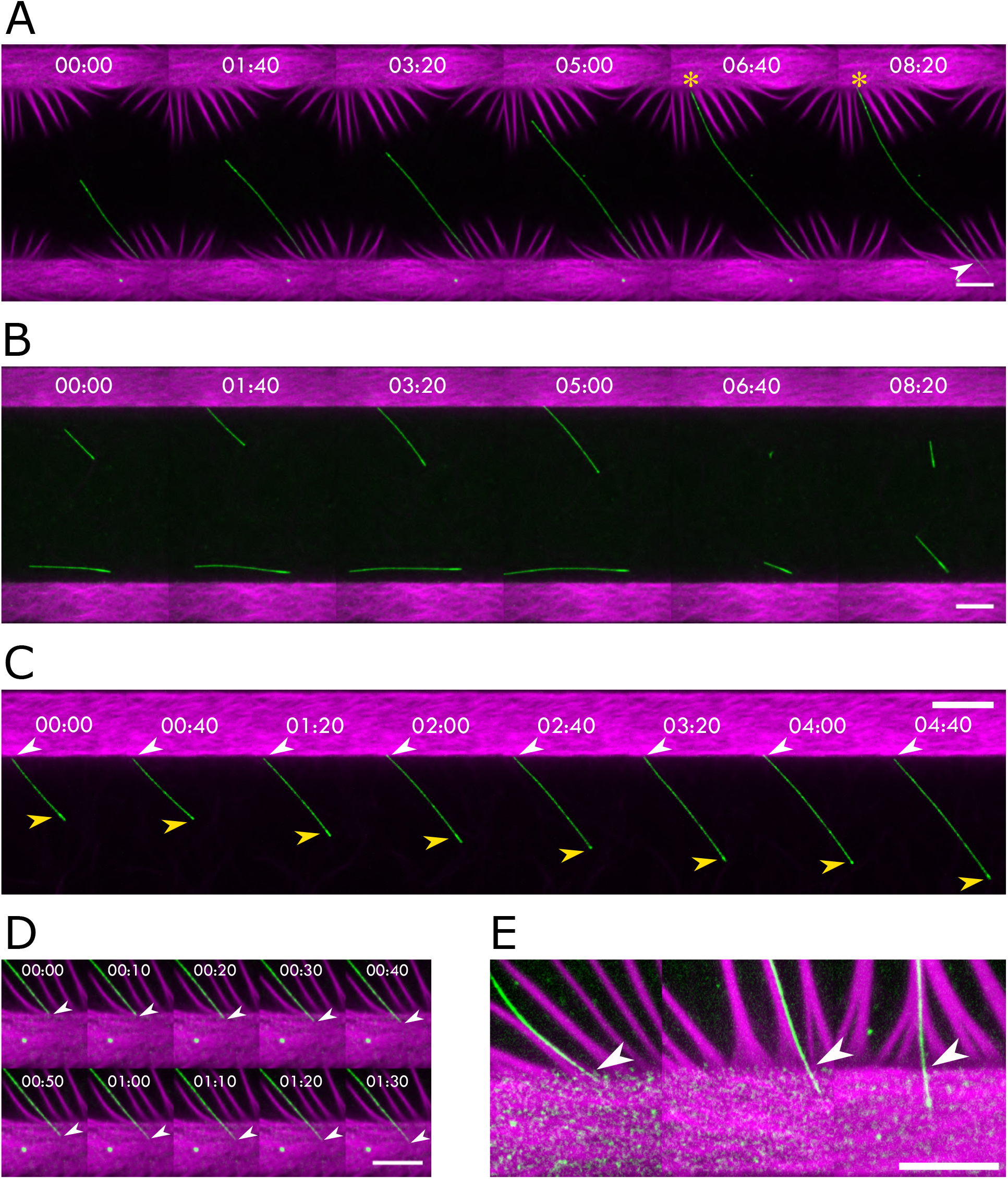
Microtubule breaching inside a dense meshwork. (**A**) Time-lapse imaging of a dynamic microtubule (green) polymerizing in between two branched actin networks (magenta) in presence of actin bundles (10nM of Arp2/3 complex). The yellow star shows the point of contact between the minus end of the microtubule and the branched network. The white arrow shows the plus end of the microtubule breaching inside of the branched actin network. (**B**) Time-lapse imaging of dynamic microtubules (green) polymerizing in between two branched actin networks (magenta) without actin bundles (100nM of Arp2/3 complex). (**C**) Zoomed time-lapse imaging of a microtubule + tip (green, white arrow) polymerizing against a branched actin network (magenta, 100nM of Arp2/3 complex). The yellow arrow indicate the microtubule seed. (**D**) Zoomed time-lapse imaging of a microtubule + tip (green, white arrow) breaching inside of the branched actin network (magenta, 10nM of Arp2/3 complex). (**E**) Snapshot of multiple microtubules + tip (green, yellow arrow) breaching inside of branched actin networks (magenta, 10nM of Arp2/3 complex). Scale bar, 10 µm.

We looked carefully at the microtubule’s growing end in contact with the actin meshwork. In the absence of actin bundle, microtubule never penetrated the actin meshwork (Figure 4C). However, in the presence of actin bundles we could see some microtubules penetrating the actin meshwork (Figure 4D). This suggested that the local organization of actin filaments at the base of the actin bundles (Svitkina et al., 2003) could be more permissive to microtubule penetration (Figure 4E). Furthermore, the alignment of microtubules with the actin bundle could facilitate this event by locking the microtubules in a favorable orientation. At the same time, we also noticed that these penetration events occurred only when both ends of the microtubules were immobilized against two adjacent actin networks (asterisk at minus-end and arrow at plus-end in Figure 4A). In such condition, microtubules could not move backward, so the polymerization force put microtubule under pressure. Noteworthy, these constrained microtubules could also pierce the meshwork from their minus end (Supplementary Figure S2). This suggested that the building up of pressure in the microtubule was responsible for their breaching through dense the actin meshwork. Such a mechanism should also promote microtubule penetration in the absence of bundles. Unfortunately, the random orientation of free microtubules in the absence of actin bundle and the distance between the two actin meshworks did not favor such event where both ends of microtubules were immobilized (Figure 4B). So we could not distinguish the role of pressure from the role of specific actin architecture associated with the formation of bundles.

### Pressure is responsible for microtubule piercing

In order to increase the number of microtubules under pressure in the absence of actin bundles, we reduced the distance between the two micropatterned bars on which actin meshwork were nucleated (Figure 5). In these conditions, we could observe the breaching of dynamic microtubules, even in the absence of actin bundles (yellow arrow head in Figure 5A). This observation shows that the specific architecture at the base of actin bundles is not necessary for microtubules to penetrate the actin network. It could nevertheless be a facilitating structure. Indeed, we observed a higher frequency of microtubules breaching the actin network in the presence of actin bundles (Figure 5B). However, in the absence of actin bundle, a significant proportion of microtubules contacted the actin network at a low angle (<30°). (Figure 5C). Because of their orientation, these microtubules were less likely to become long to make contact with the two branching networks at both ends (Figure 5A, bottom). To compare microtubules with similar probability to be pressurized by their growth, we selected only the microtubules that contacted the actin meshwork with an angle higher than 30° (Figure 5D). Then, there was no more difference in the breaching probability of microtubule in the presence or absence of bundle of actin filament. This showed that the presence of actin bundle did not seem to facilitate the penetration of microtubule in the actin meshwork, and that the increase of pressure in the microtubule was the main parameter responsible for these events.

**Figure 5.**
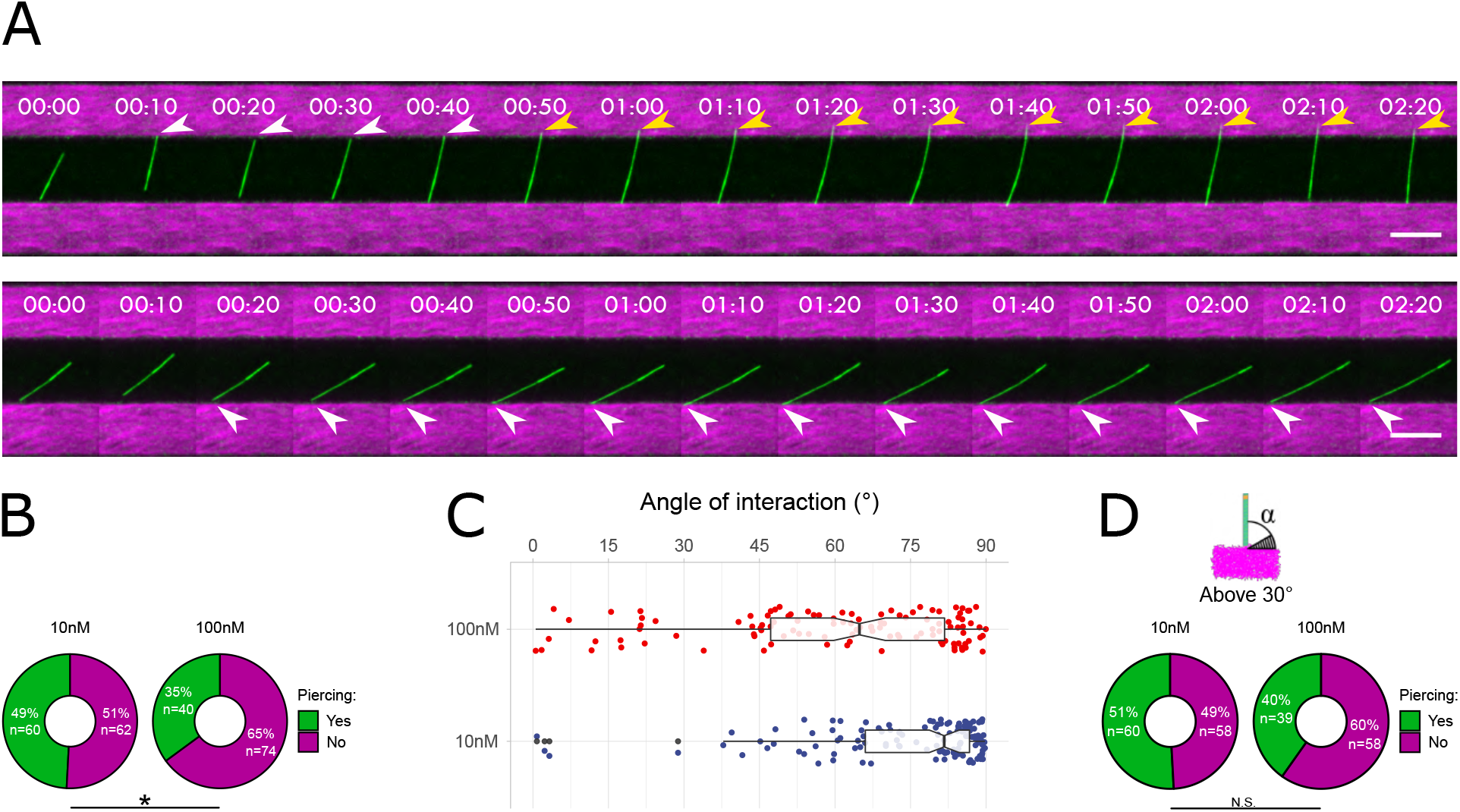
Resisting load favor piercing. (**A**) Time-lapse imaging of dynamic microtubules (green) polymerizing in between two branched actin networks (magenta), without actin bundles (100nM of Arp2/3 complex), at an angle lower (bottom) or greater than 30°(top). White arrows show the point of contact between microtubules + tip and the branched actin meshwork. Yellow arrows show microtubule + tip piercing the branched actin meshwork. (**B**) Distribution of the angle of interaction of microtubules in the presence (10nM of Arp2/3 complex) or absence (100nM of Arp2/3 complex) of actin bundles. (**C**) Proportion of microtubules piercing the branched actin network in two different conditions: 10nM of Arp2/3 complex and 100nM of Arp2/3 complex. * p<0.05 (Chi-squared test χ2). (**D**) Proportion of microtubules piercing the branched actin network in two different conditions: 10nM of Arp2/3 complex and 100nM of Arp2/3 complex. Only the microtubules with an interaction angle superior to 30°were included in the analysis. N.S. p>0.05 (Chi-squared test χ2). Scale bar, 10 µm.

### Crosslinking between microtubules and actin bundles increase piercing frequency

In cells, microtubules can penetrate the lamellipodium. The local buckling of microtubule shows that they are not free to move backward as they grow against the lamellipodium (Wittmann et al., 2003; Gupton et al., 2002) and that they are likely attached to some structures, which induce the build-up of pressure at the growing end. Indeed, microtubules are tethered to actin bundles or to other microtubules via specific crosslinkers (Rodriguez et al., 2003; Bodakuntla et al., 2019). We thus tested whether such crosslinkers could contribute to the build-up of pressure and to the consequent breaching of microtubules through dense actin meshwork in our in vitro assay. To that end, we selected the protein Tau to crosslink microtubules with actin bundles (Elie et al., 2015; Cabrales Fontela et al., 2017) and allow the transmission of polymerization forces between the two (Alkemade et al., 2022). We worked in the 10nM of Arp2/3 complex condition to allow actin bundles to grow out of the dense meshwork and added 100nM of Tau to crosslink them to dynamic microtubules (Figure 6). As expected, in the control condition, the microtubules nicely align with the actin bundles, perpendicular to the branched network, and move rearward due to the force produced by their polymerization (Figure 6A). They rarely penetrate the actin network (6%, 3 out of 47 microtubules, Figure 6B). By contrast, in the presence of Tau, microtubules could not move backward and we could detect clear beaching events (Figure 6C). The frequency of these breaching events was dramatically increased by the presence of Tau (40%, 31 out of 77 microtubules, Figure 6B). In these conditions, we cannot exclude that Tau also affected the architecture of the actin network (Elie et al., 2015). However, these results strongly suggest that the binding of microtubules to actin bundles build up some pressure in the growing microtubule, which promoted its penetration in the densely branched actin meshwork.

**Figure 6.**
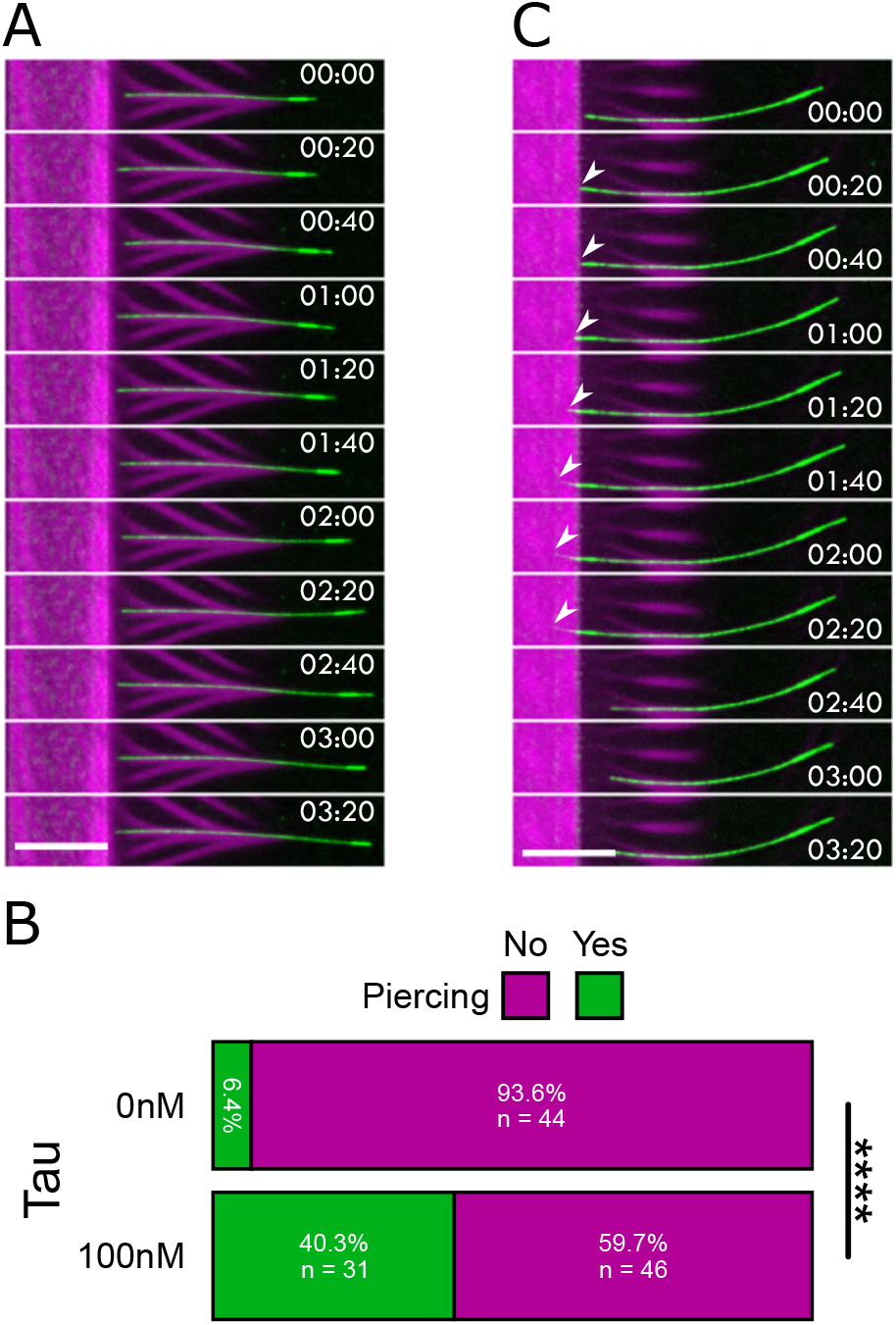
The presence of crosslinker favors piercing. (**A**) Time-lapse imaging of dynamic microtubules (green) polymerizing against a branched actin network (magenta) in presence of actin bundles (10nM of Arp2/3 complex). (**B**) Proportion of microtubules piercing the branched actin network in two different conditions: 0nM of Tau and 100nM of Tau. **** p<0.0001 (Chi-squared test χ2). (**C**) Time-lapse imaging of dynamic microtubules (green) polymerizing against a branched actin network (magenta) in presence of actin bundles (10nM of Arp2/3 complex) and of an actin-microtubule crosslinker (100nM of Tau). White arrows show microtubule + tip piercing the branched actin meshwork. Scale bar, 10 µm.

## Discussion

Altogether, our results highlight the role of structural components, which, by resisting the backward translocation of growing microtubules, induce the build-up of pressure that push them to breach into dense actin meshworks. Our results also showed that if actin and microtubules are free to move, they will tend to align with each other even in the absence of crosslinkers. This alignment could be due to electrostatic or hydrophobic interactions (Schaedel et al., 2021). It could also result from the presence of methyl-cellulose in our polymerization mix, since crowding agents promote the alignment of long structures through colloid osmotic pressure (Mitchison, 2019). It was tempting to speculate that this alignment guide microtubules toward regions where the organizing of actin filaments is more permissive to penetration (Schaefer et al., 2002), but, although we could not fully exclude this hypothesis, our results did not provide any evidence for such a scenario.

Whether the density of branched actin meshwork in our reconstitution assay is qualitatively comparable to the lamellipodium meshwork in cells is hard to tell. However, it should be noted that the frequency of microtubule penetrating is low in both conditions, suggesting that the conditions are likely similar. Interestingly, our results show that the forces produced by microtubule polymerization are sufficient to pierce the said dense and branched actin meshwork but that counter forces are required for this to happen. This means that in cells, the frequency of these events is under the control of the structures that can bind to microtubules and resist their backward translocation upon elongation. This suggest that the local regulation of the concentration of molecular motors, crosslinkers, or others means, might allow microtubules to breach through the lamellipodium in some specific places and thereby control the location of the delivery of specific cargos transported along microtubules. This capacity to locally adjust signals delivery might be instrumental in the context of cell shape remodeling and migration.

## Methods

### Cell culture

#### PtK_2_

Cell culture Male rat kangaroo kidney epithelial cells (PtK_2_) stably expressing GFP-Tubulin obtained from Franck Perez lab were grown at 37°C and 5% CO2 in DMEM/F12 (31331028, Gibco) supplemented with 10% fetal bovine serum (10270106, Life Technologies) and 1% antibiotic-antimycotic solution (15240062, Gibco).

#### MEF

Wild type mouse embryonic fibroblasts cells (MEF WT) obtained from the lab of John Eriksson were grown at 37°C and 5% CO2 in DMEM/F12 (31331028, Gibco) supplemented with 10% fetal bovine serum (10270106, Life Technologies) and 1% antibiotic-antimycotic solution (15240062, Gibco)

#### COS-7

Cells were cultured in DMEM high glucose/Glutamax supplemented with 10% fetal bovine serum (Thermo Fisher). Fixation and staining was performed according to a previously published protocol (Jimenez et al., 2020). Briefly, cells were extracted in Triton X-100/glutaraldehyde, then fixed using glutaraldehyde, before being quenched sing NaBH4, blocked and stained with anti-alpha tubulin primary antibodies (monoclonal mouse clones B-5-1-2 and DM1a, Sigma) overnight at 4°C, which were revealed with donkey antimouse secondary antibodies conjugated to Alexa Fluor 555 (1h at RT). To stain actin, cells were incubated for 1h with phalloidin-Atto488 (Sigma) at the end of the staining procedure, and mounted in Prolong Glass medium (Thermo Fisher).

### Cell fixation and labeling of actin and microtubules

Prior to fixation MEF cells were plated overnight on glass coverslips coated with fibronectin at 10 µg/mL. Cells were fixed 10 min at RT in cytoskeleton buffer (MES 10mM, KCl 138mM, MgCl_2_ 3mM, EGTA 2mM) supplemented with 10% sucrose, 0.05% Triton-X100, 0.25% glutaraldehyde (G5882, Sigma), and 4% of paraformaldehyde. Aldehyde functions were then reduced using 1mg/ml of NaBH4 for 10 minutes at RT. The samples were then washed 3 times with PBS-Tween 20 (1379, Sigma) 0.1% and incubated in a blocking solution (PBS, 0.1% Tween-20, 3% BSA (A2153, Sigma)) for 25 min at RT. This was followed by an incubation in blocking solution at RT for 30 min with a primary antibody directed against tyrosinated tubulin (YL1/2 : MAB1864, Merk) used at 1/400. The slides were then rinsed 3 times using PBS-Tween 0.1% and incubated in blocking solution for 30 min at RT with the secondary antibody (712-545-153, Jackson Immunoresearch) at 1/400, DAPI (D9542, Sigma) at 1/1000, and phalloidin (A34055, Life Technologies) at 1/400. Finally, the slides were rinsed twice in PBS-Tween 0.1%, once in PBS and mounted in Mowiol 4-88 (81381, Sigma).

### Lipids/SUV preparation

Clean the glass tube with water, EtOH and chloroform. Dry it using azote. Make the lipids mix (98.75% EggPC (Avanti, 840051C) 10mg/mL, 0.25% PEG-biotin 10mg/mL, 1% DOPE-Atto 390 1mg/mL) using Hamilton syringe cleaned with chloroform. Dry with azote on vortex under the hood. Put under vacuum overnight. Resuspend with 1mL of SUV buffer-vortex (10mM Tris pH 7.4, 150mM NaCl, 2mM CaCl_2_). The mixture was sonicated on ice for 10 min. The mixture was then centrifuged for 10 min at 20,238 g to remove large structures. The supernatants were collected and stored at 4°C.

### Protein purification

#### Tubulin

Bovine brain tubulin was purified in BRB80 buffer (80 mM 1,4-piperazinediethanesulfonic acid, pH 6.8, 1 mM ethylene glycol tetraacetic acid, and 1mM MgCl_2_) according to previously published method (Vantard et al., 1994). Tubulin was purified from fresh bovine brain by three cycles of temperature-dependent assembly and disassembly in Brinkley Buffer 80 (BRB80 buffer; 80 mM PIPES, pH 6.8, 1 mM EGTA, and 1 mM MgCl_2_ plus 1 mM GTP) (Shelanski, 1973). MAP-free neurotubulin was purified by cation-exchange chromatography (EMD SO, 650 M, Merck) in 50 mM PIPES, pH 6.8, supplemented with 1 mM MgCl_2_, and 1 mM EGTA. Purified tubulin was obtained after a cycle of polymerization and depolymerization. Fluorescent tubulin (ATTO 488–labeled tubulin and ATTO 565–labeled tubulin) and biotinylated tubulin were prepared according to previously published method Hyman et al. (1991). Microtubules from neurotubulin were polymerized at 37°C for 30 min and layered onto cushions of 0.1 M NaHEPES, pH 8.6, 1 mM MgCl_2_, 1 mM EGTA, 60% v/v glycerol, and sedimented by high centrifugation at 30°C. Then microtubules were resuspended in 0.1 M NaHEPES, pH 8.6, 1 mM MgCl_2_, 1 mM EGTA, 40% v/v glycerol and labeled by adding 1/10 volume 100 mM NHS-ATTO (ATTO Tec), or NHSBiotin (Pierce) for 10 min at 37°C. The labeling reaction was stopped using 2 volumes of 2X BRB80, containing 100 mM potassium glutamate and 40% v/v glycerol, and then microtubules were sedimented onto cushions of BRB80 supplemented with 60% glycerol. Our labelling ratio is: for tubulin 488 nm 1.6 to 1.8 fluorophore by dimers and for tubulin 565 nm 0.6 to 0.7 fluorophore by dimers. We used NanoDrop one UV-Vis spectrophotometer (Thermofisher) to determine the labelling efficiency. Microtubules were resuspended in cold BRB80. Microtubules were then depolymerised and a second cycle of polymerization and depolymerization was performed before use.

#### Actin

Actin was purified from rabbit skeletal-muscle acetone powder (Spudich and Watt, 1971). Monomeric Ca-ATP-actin was purified by gel-filtration chromatography on Sephacryl S-300 at 4°C in G-Buffer (5 mM Tris-HCl (pH 8.0), 0.2 mM ATP, 0.1 mM CaCl_2_ and 0.5 mM dithiothreitol (DTT)). Two grams of muscle acetone powder were suspended in 40 mL of buffer G and extracted with stirring at 4°C for 30 min, then centrifuged 30 min at 30,000g at 4°C. The supernatant with actin monomers was filtered through glass wool and we measured the volume. The pellets were suspended in the original volume of G-Buffer and we repeated the centrifugation and filtration steps. While stirring the combined supernatants in a beaker add KCl to a final concentration of 50mM and then 2 mM MgCl_2_ to a final concentration of 2 mM. This step will polymerize the actin monomers. After 1 h, add KCl to a final concentration of 0.8 M while stirring in cold room. This dissociates any contaminating tropomyosin from the actin filaments. After 30 min, centrifuge 2 h at 140,000g to pellet the actin filaments. Discard supernatant and gently wash off the surface of the pellets with G-buffer. Gently suspend the pellets in about 3 mL of G-buffer per original gram of acetone powder using a Dounce homogenizer and dialyze for 2 days vs. three changes of G-Buffer to depolymerize the actin filaments. To speed up depolymerization, you can sonicate the suspended actin filaments gently. Clarify the depolymerized actin solution by centrifugation in Ti45 rotor at 140,000Ög for 2 h to remove aggregates. The top 2/3 of the ultracentrifuge tube contains “conventional” actin. Gel filter on Spectral S-300 in buffer G to separate actin oligomers. Labeling was done on lysines by incubating actin filaments with Alexa-568 succimidyl ester (Molecular Probes).

#### Arp2/3 complex

The Arp2/3 complex was purified from calf thymus according to (Egile et al., 1999) with the following modifications: the calf thymus was first mixed in an extraction buffer (20 mM Tris pH 7.5, 25 mM KCl, 1 mM MgCl_2_, 0.5 mM EDTA, 5% glycerol, 1 mM DTT, 0.2 mM ATP and proteases). Then, it was placed in a 50% ammonium sulfate solution in order to make the proteins precipitate. The pellet was resuspended in extraction buffer and dialyzed overnight. The Arp2/3 complex was fluorescently labeled as described in Funk et al. (2021).

#### Streptavidin-WA

Snap-Streptavidin-WA-His (pETplasmid) was expressed in Rosetta 2 (DE3) pLysS (Merck, 71403). Culture was grown in TB medium supplemented with 30 µg/ml kanamycin and 34 µg/ml chloramphenicol, then0.5 mM isopropyl β-D-1-thiogalactopyranoside (IPTG) was added, and protein was expressed overnight at 16°C. Pelleted cells were resuspended in Lysis buffer (20 mM Tris pH8, 500 mM NaCl, 1 mM EDTA, 15 mM Imidazole, 0.1% TritonX100, 5% Glycerol, 1 mM DTT). Following sonication and centrifugation, the clarified extract was loaded on a Ni Sepharose high-performance column (GE Healthcare Life Sciences, ref 17526802). Resin was washed with Wash buffer (20 mM Tris pH8, 500 mM NaCl, 1 mM EDTA, 30 mM Imidazole, 1 mM DTT). Protein was eluted with Elution buffer (20 mM Tris pH8, 500 mM NaCl, 1 mM EDTA, 300 mM Imidazole, 1 mM DTT). Purified protein was dialyzed overnight 4°C with storage buffer (20 mM Tris pH8, 150 mM NaCl, 1 mM EDTA, 1 mM DTT), concentrated with Amicon 3KD (Merck, ref UFC900324) to obtain concentration around 10 µM then centrifuged at 160,000 g for 30 min. Aliquots were flash-frozen in liquid nitrogen and stored at -80°C.

### Silane-PEG slides and coverslip

The micropatterning technique was adapted from (Portran et al., 2013). Cover glasses were cleaned by successive chemical treatments: 30 min in acetone, 15 min in ethanol (96%), rinsing in ultrapure water, 2 h in Hellmanex III (2% in water, Hellmanex), and rinsing in ultrapure water. Cover glasses were dried using filtered air flow, oxidized using air plasma during 5min (400 W, PlasmaEtch), and incubated at least overnight in a solution of tri-ethoxysilane-PEG (5kDa for coverslip and 30 kDa for slides, Creative PEGWorks) at 1mg *\** ml–1 in ethanol 96% and 0.02% HCl, with gentle agitation at room temperature. Cover glasses were kept in solution at room temperature and stored until use. When used, the cover glasses were successively washed in ethanol, ultrapure water and dried with filtered air.

### PRIMO patterning

A chamber made of a passivated coverslip and cover glass were mounted using some double-sided tape. This chmaber was then perfused with a photoinitiator solution (PLPP, Alveole) and transferred on an inverted microscope (Ti-E, Nikon) equipped with the PRIMO system (Alveole). To create patterns on silane-PEG a UV dose of 100mJ/mm^2^ was used. Once the UV exposure was done the chamber was rinsed using a SUV buffer (10mM Tris pH 7.4, 150mM NaCl, 2mM CaCl_2_). Prior to the photopatterning, the alignment of the motorized stage with the camera and calibration of the Digital Micromirror Device (DMD) with the camera was done using the Leonardo software (Alveole) (Strale et al., 2016).

### Microtubule and patterned actin interaction assay

Microtubule and actin networks were reconstituted In vitro. A flow cell with an approximate reaction volume of 6*µ*L with an entry and exit sites were constructed using a double-sided tape (70*µ*m height) between a glass coverslip coated with SiPEG 5kDa, a passivated glass slide coated with SiPEG 30 kDa, and patterned using the PRIMO system as described in the previous sections. After the patterning, a first washing step was done using the SUV buffer (10mM Tris pH 7.4, 150mM NaCl, 2mM CaCl_2_) in order to remove the excess PLPP. Next, the lipids were introduced in the chamber and incubated for 3min, followed by a washing step with the SUV buffer and another washing using the “wash buffer” solution composed of a mix of 1x HKEM (6.7mM HEPES pH 7.5, 50mM KCl, 5mM MgCl_2_, 1mM EGTA) and 1x G-buffer (2mM Tris-HCl (pH 8.0), 0.2mM ATP, 0.1mM CaCl_2_ and 0.5mM dithiothreitol (DTT)). Next, a dilution of streptavidin-WA (100nM in wash buffer) was incubated for 3min. A final rinse was done using the washing buffer. Finally, the solution of “Tic-Tac” buffer was introduced containing the actin all its necessary related proteins and tubulin: Actin (1*µ*M), The Arp2/3 complex (between 10nM or 100nM), Human Profilin (between 2 times the actin concentration), Tubulin (17*µ*M), Microtubule seeds (1:750 dilution in wash buffer) and Tau (100nM).

### Microscopic observation

#### TIRF imaging

Imaging of experiments involving microtubules were performed on an inverted microscope (Ti-E, Nikon), equipped with a total internal reflection fluorescence (TIRF) iLasPulsed system (Gataca Systems) and a Retiga R3 camera (CCD 1920×1460, Binning = 1, pixel = 4.54*µ*m) using an Olympus U APO N TIRF oil-immersion 100× 1.49 N.A. objective lens, with a 0.63 O ring. The microscope stage was maintained at 37°C by means of a temperature controller to obtain an optimal microtubule growth. Multi-stage time-lapse movies were acquired using Metamorph software (version, Universal Imaging).

For experiments involving the observation of big pattern over the course of an experiment, a custom stage list was created using the scan slide journal of Metamorph and the origin of the stage was setup as the top right corner of the pattern.

### Imaging of fixed samples

Acquisition of fixed samples were made with an inverted microscope (Ti-E, Nikon), equipped with a spinning disk confocal csu (Yokogawa CSU-X1), iLasPulsed system (Gataca Systems) and a Retiga R3 camera (CCD 1920×1460, Binning = 2, pixel = 4.54µm) using an Nikon plan APO VC oil-immersion 60x 1.40 N.A. Multistage Z series acquisition were done using Metamorph software (version 7.7.11.0, Universal Imaging).

#### Live imaging of actin and microtubules in cells

PtK_2_ cells were plated overnight in glass bottom dishes (627860, Dutscher) coated with fibronectin and collagen both at 10 µg/mL. Cells were then treated for 4h with SiR-Actin (SC001, tebubio) at concentrations between 200 and 300 nM and Verapamil (SC001, tebubio) at concentrations between 10 and 15µM. Live acquisitions of endogenously expressed GFP tubulin and filamentous actin stained with SIR-Actin were made inside a live module (kept at 37°C and 5% CO2) mounted on a confocal spinning disk microscope (Nikon Ti Eclipse equipped with a spinning scanning unit CSU-X1 Yokogawa) and a R3 retiga camera (QImaging). Images were acquired using a Nikon Plan Apo VC 60x/1.40 NA oil objective. Each wavelength was acquired separately. Metamorph software was used for images acquisition (version 7.7.11.0, Universal Imaging).

#### SIM imaging

Cells were imaged on a N-SIM-S microscope (Nikon) using a classical 3-beam configuration for 3D-SIM. Laser at 488 nm (for actin) and 561 nm (for microtubules) were used to illuminate the sample through a 100X, 1.49 NA objective, capturing 15 images (16-bit, 1024×1024 pixels at 65 nm/pixel) over a 50 to 100 ms exposure time via a Fusion BT sCMOS camera (Hamamatsu). The raw images were then processed using the N-SIM module of the NIS Elements software, resulting in a 32-bit, 2048×2048 pixel reconstructed image at 32.5 nm/pixel. Images were converted to 16 bit, projected, overlayed and contrast-adjusted in Fiji.

### Statistical analysis

All the graphs and statistical analysis were performed using R (Core Team R, 2023).

## Acknowledgement

We would like to thank the Neuro-Cellular Imaging Service and Nikon Center for Neuro-NanoImaging at INP, with funding from CPER-FEDER (PlateForme NeuroTimone PA0014842) as well as the Institut Marseille Imaging for complementary equipment funding from Excellence Initiative of Aix-Marseille University -A*MIDEX, a french “Investissements d’Avenir” programme (AMX-19-IET-002).

## Supplementary Information

**Figure S1.**
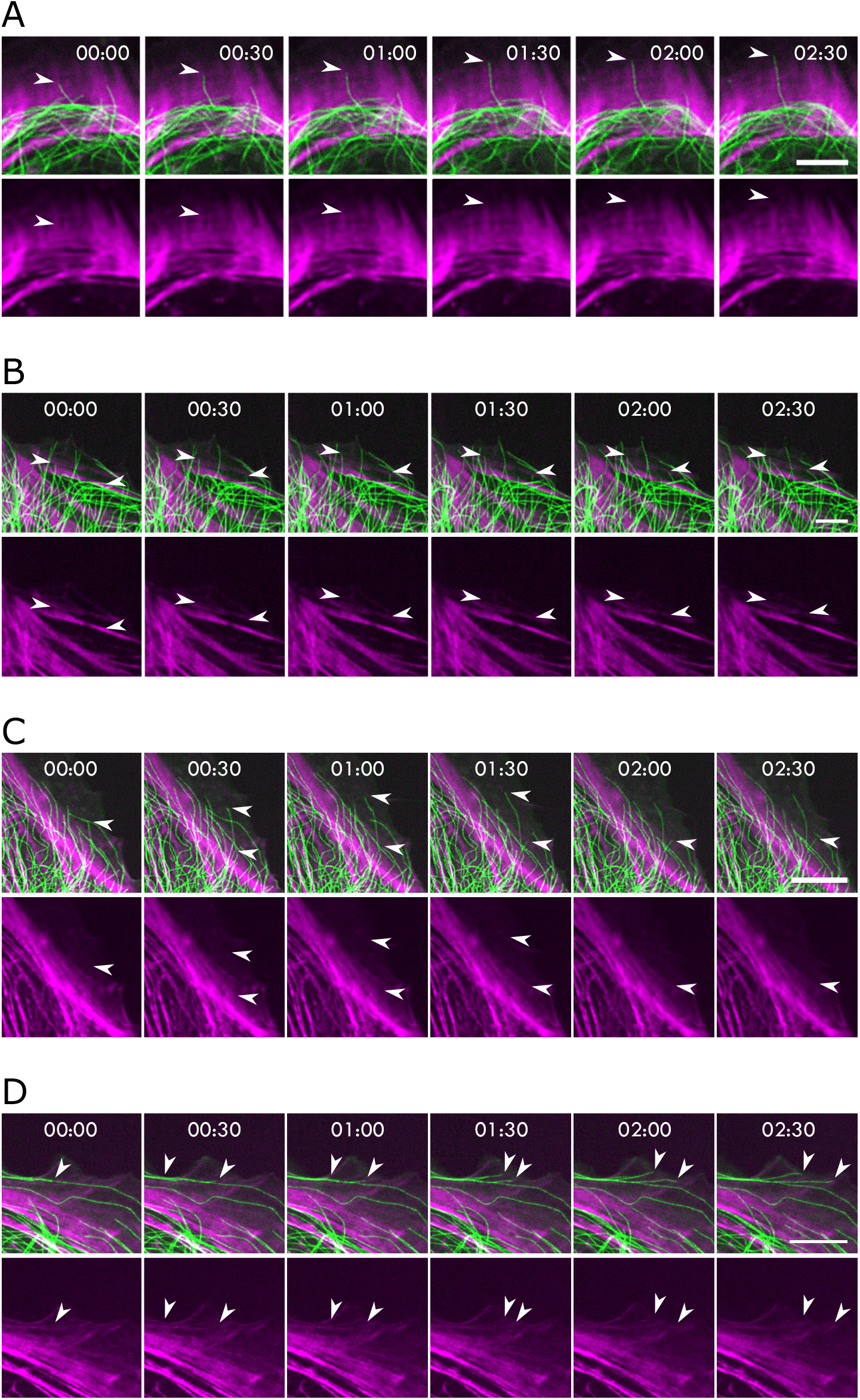
Examples of microtubules entering the lamellipodial region of cells. (**A-B-C-D**) Time-lapses of a PtK_2_ cell tubulin-GFP (green) incubated with 200nM of SiR-actin (magenta). The white arrows are showing microtubules entering the lamellipodial region of the cell (Top) and the actin structure encountered along their path (bottom). Scale bar, (**A-B**) 5 µm, (**C-D**) 10 µm

**Figure S2.**
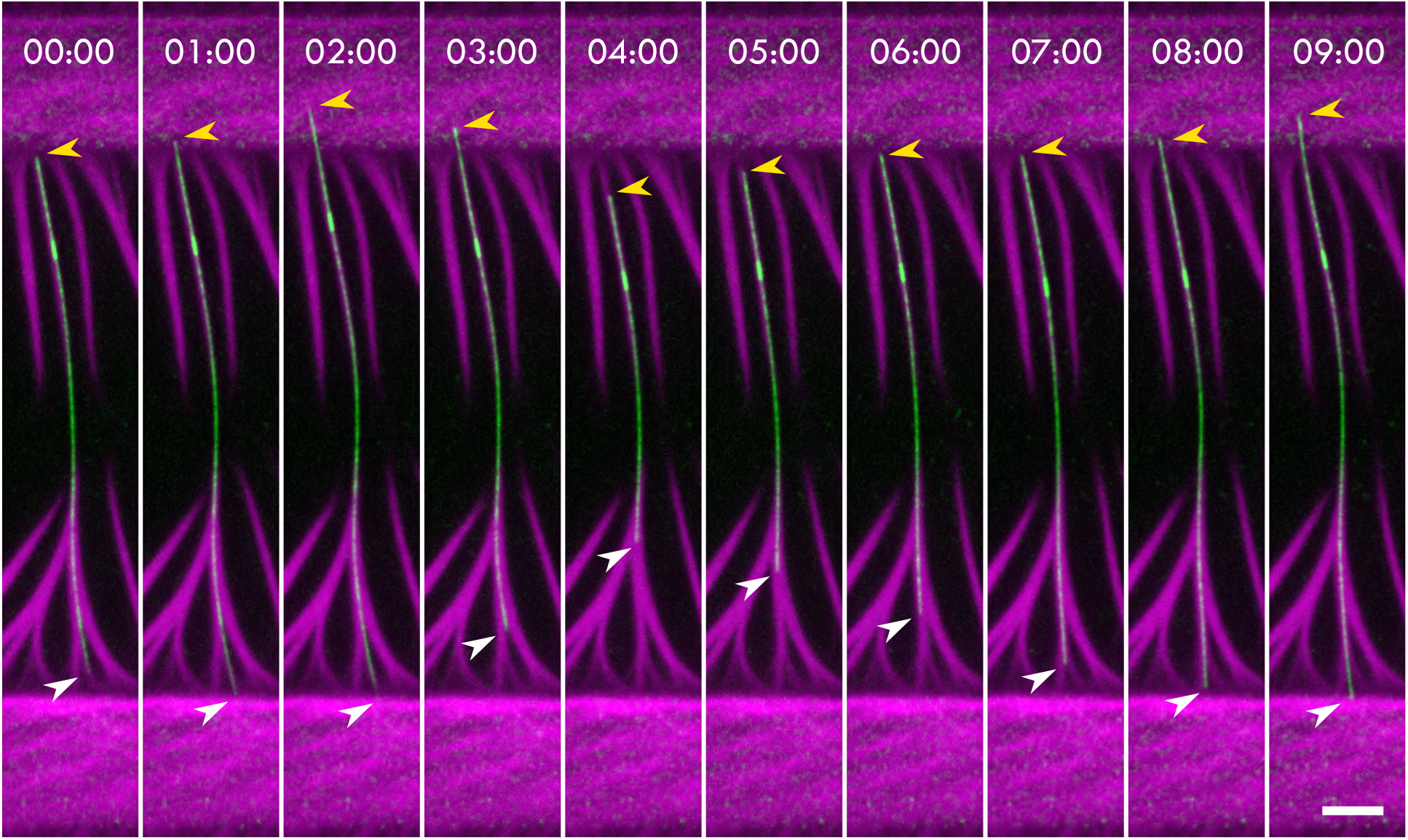
Microtubule piercing from the minus end. Time-lapse of a microtubule + tip (green, white arrow) polymerizing against a branched actin network (magenta, 10nM of Arp2/3 complex). The yellow arrow indicate the microtubule -tip. Under the pressure of polymerization the minus tip of the microtubule is able to breach inside the branched actin network. Scale bar, 5 µm.

